# Multifunctional molecular hybrid for targeted colorectal cancer cells: Integrating doxorubicin, AS1411 aptamer, and T9/U4 ASO

**DOI:** 10.1101/2024.05.08.593145

**Authors:** Kanpitcha Jiramitmongkon, Pichayanoot Rotkrua, Paisan Khanchaitit, Jiraporn Arunpanichlert, Boonchoy Soontornworajit

## Abstract

Colorectal cancer (CRC) is one of the public-health concerns worldwide and it requires an effective treatment. However, existing treatment approaches encounter challenges related to specificity and efficacy. To address this issue, a platform for multifunctional drug delivery has been developed, combining bioactive materials with anticancer elements and specific recognition ligands into a single molecule. This study aimed to create a molecular hybrid (MH) containing doxorubicin, AS1411 aptamer, and T9/U4 ASO to regulate SW480 cell proliferation. The AS1411 aptamer targets nucleolin, overexpressed on cancer cell membranes, while T9/U4 ASO inhibits human telomerase RNA activity, further hindering cancer cell proliferation. AS-T9/U4_MH was synthesized via oligonucleotide hybridization, followed by doxorubicin loading and evaluation of its impact on cell proliferation. Binding capability of this MH was verified using fluorescence microscopy and flow cytometry, demonstrating specific recognition of SW480 cells due to nucleolin availability on the cell surface. These findings were corroborated by both microscopy and flow cytometry. AS-T9/U4_MH exhibited anti-proliferative effects, with the doxorubicin-loaded system demonstrating encapsulation and reduced toxicity. Moreover, the presence of Dox within AS-T9/U4_MH led to a notable reduction in hTERT and vimentin expression in SW480 cells. Additionally, examination of apoptotic pathways unveiled a marked decrease in Bcl-2 expression and a simultaneous increase in Bax expression in SW480 cells treated with Dox-loaded AS-T9/U4_MH, indicating its impact on promoting apoptosis. These results suggest that the molecular hybrid holds promise as a system for integrating chemotherapeutic drugs with bioactive materials for cancer treatment delivery.

## Introduction

The progression of cancer involves intricate mechanisms that engage multiple biomolecules [1]. Aiming to enhance the efficacy of anti-cancer drugs, the concept of multifunctional drugs has emerged. These drugs typically exhibit theranostic characteristics, simultaneously serving as diagnostic probes and specific drug delivery vehicles to targets. This dual functionality ensures efficient performance [2]. Moreover, multifunctional drugs can enhance treatment efficiency by increasing their accumulation in target tissues while minimizing drug availability in other organs, thereby reducing adverse effects [3]. Literature reports reveal the integration of a redox-responsive hyperbranched polymer with a nucleolin aptamer and doxorubicin to create a multifunctional drug delivery system. The aptamer acts as both a recognition element and a biological probe, while doxorubicin serves as the therapeutic agent for cancer. This innovative system has proven effective in mitigating severe side effects associated with cancer treatment and enhancing overall treatment efficiency [4]. The formation of multifunctional drugs employs a combination of noncovalent interactions and chemical strategies. These approaches have been successfully implemented in various forms, including hybrid nanogels, hollow microspheres, and wearable transdermal delivery systems [5–7]. The outcomes of these strategies include soft-matter nanoarchitectures with defined sizes and morphologies, tunable luminescence, and specific biological functions [2]. One notable strategy involves the hybridization of complementary oligonucleotides to form double-stranded DNA, a facile approach for creating complex structures in multifunctional drugs [8]. This method allows for the integration of functional oligonucleotides such as antisense oligonucleotides (ASOs), small interfering RNAs (siRNAs), and therapeutic aptamers [9, 10]. The current work proposes to expand on our understanding of multifunctional drug delivery systems, particularly those comprising aptamers and ASOs, by providing additional information and insights.

Aptamers are unique DNA or RNA sequences composed of a single strand, identified through the systematic evolution of ligands by exponential enrichment (SELEX) process [11]. This method produces sequences characterized by exceptional binding capabilities, displaying both high specificity and affinity. Additionally, aptamers possess favorable attributes such as ease of chemical modification, low immunogenicity, commercial availability, and a broad range of target options [11]. The AS1411 aptamer, specifically, is a short single-stranded DNA with a 26-mer G-rich region. It serves as a recognition ligand for nucleolin, which is overexpressed on the surface of numerous cancer cells with remarkable selectivity and affinity. Moreover, the AS1411 aptamer, adopting a G-quadruplex conformation, has demonstrated anti-proliferation activity [12], increased resistant to nucleases [13] and enhanced cellular uptake [14]. Consequently, the AS1411 aptamer emerges as a promising candidate for use as a carrier for antisense oligonucleotides targeted at specific cancer cells.

ASOs are large nucleic acid molecules that selectively regulate the functions of a target gene. By virtue of their complementary nature to target mRNA, they engage in Watson-Crick base pairing, leading to the destabilization and degradation of the mRNA and, consequently, interference with the translation of the associated protein [15]. ASOs typically operate through two primary modes of action: splicing modulation and gene expression inhibition. In the role of splicing modulators, ASOs bind specifically to mRNA regions, blocking ribosome binding and thereby suppressing protein translation. As gene expression modulators, ASOs form duplexes with RNA, prompting RNAse H to degrade the targeted RNA sequences [16]. Numerous ASOs have advanced into clinical trials and gained approval for therapeutic use, such as in cases like VITRAVENE^®^ (fomivirsen), Kynamro^®^ (mipomersen), and Spinraza^®^ (nusinersen) [17]. These ASOs exhibit low cytotoxicity, generating substantial interest in the development of effective anticancer formulations [18]. In existing literature, a specific ASO sequence known as T9/U4 has shown promising anticancer potential by inhibiting telomerase RNA [19]. Telomerase, a reverse transcriptase enzyme, comprises an RNA component (human telomerase RNA (hTR) or human telomerase RNA component (hTERC)) and a protein subunit (human telomerase reverse transcriptase (hTERT)) [20]. The proposed mechanism of action for this ASO involves direct inhibition of enzymes at the hTR active site, targeting the catalytic dysfunction of hTERT [21].

To suppress gene expression, the penetration of ASOs into target cells is essential. However, the precise mechanisms remain unclear, potentially involving factors such as size [22], structure [23], concentration, and cell line [24]. Challenges associated with ASO utilization include limited stability, non-specific delivery to target cells, and low cellular uptake, impeding their silencing effect [25]. Various delivery strategies, including viral vectors [26] and non-viral vectors [27] like liposomes, polymeric, and metal nanoparticles [28], have been developed to address these issues. Nevertheless, some of these systems are effective in cultured cells but face limitations in vivo biodistribution. Additionally, cationic lipids and polymers may induce adverse effects [29], while metal nanoparticles could accumulate in cells and tissues, leading to mutation [30]. To overcome these challenges, molecular hybridization techniques have emerged as promising approaches for ASO and aptamer delivery. This method involves using ASOs with complementary sequences, exhibiting bioactivity, such as aptamers, to form double-stranded DNA and facilitate accumulation in target cells. Nucleic acid aptamers prove advantageous as intracellular delivery vehicles due to their specific and strong binding to cell surface receptor molecules [31]. Designed for versatile applications, these aptamers can be coupled with nanoparticles [32], drugs [33], and other nucleic acids [34]. Aptamers, serving as carriers for antisense oligonucleotides (ASOs), were developed with the goals of minimizing ASO dosage, mitigating off-target effects, enhancing cytocompatibility, improving selectivity, and optimizing cellular uptake [35].

An essential element in chemotherapy is the use of anticancer drugs. However, many drugs employed in treatment lack precision in distinguishing between diseased cells and normal cells, impacting not only fast-growing pathological cells but also impeding the growth of healthy cells. Therefore, there is a crucial need for the development of precise therapeutic agents. Doxorubicin (Dox), an anthracycline drug, is commonly utilized in treating various cancers, such as gastric [36], lung [37], and ovarian cancer [38]. Despite its effectiveness, Dox has limitations due to side effects such as cardiotoxicity, nausea, vomiting, fatigue, alopecia, and oral sores [39]. To mitigate these adverse effects, Dox is often used in conjunction with other medicinal herbs [40] and drugs [41] to reduce toxicity, especially in cardiac tissues. Notably, Dox’s structural composition includes flat aromatic moieties that intercalate between DNA base pairings making it a suitable candidate for incorporation into molecular hybrids [42]. In this study, a molecular hybrid named Dox-loaded AS-T9/U4_MH, comprising doxorubicin, the AS1411 aptamer, and the T9/U4 antisense oligonucleotide (ASO), was prepared, characterized, and assessed for its anti-cancer activities (Scheme 1). The investigation focused on the primary adenocarcinoma colorectal cancer cell lines SW480 and Caco-2, along with the human normal colon cell line CCD 841 CoN, to evaluate the activities of the proposed molecular hybrid.

**Scheme 1.**
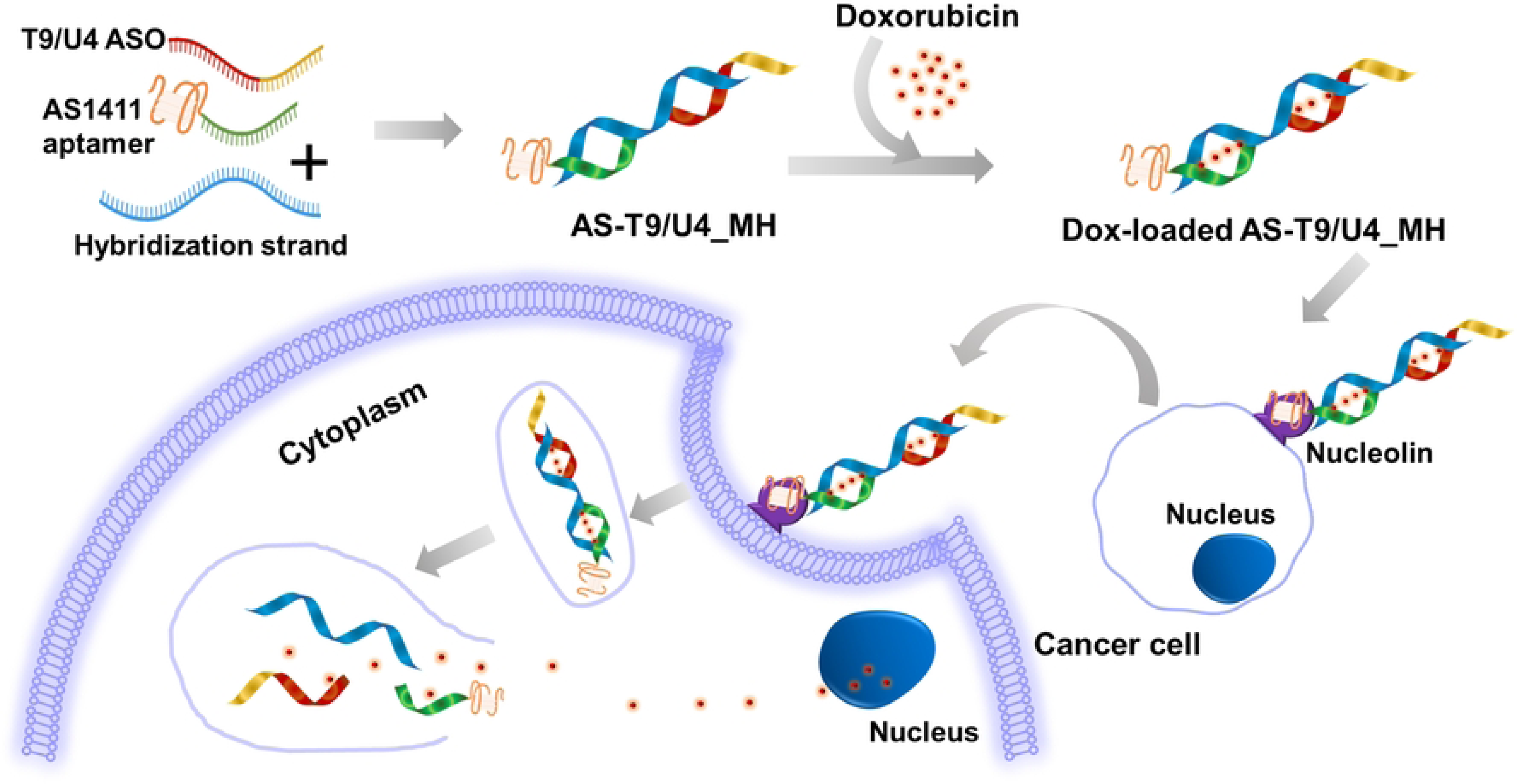
The concept for preparing Dox-loaded AS-T9/U4_MH.

## Methods

### Reagent

All the DNA molecules listed in Table 1 were purchased from Integrated DNA Technologies (IDT). Acrylamide/bis-acrylamide, tris-borate-EDTA buffer, ammonium persulfate, and gel loading buffer were obtained from Sigma-Aldrich. *N,N,N′,*N′-Tetramethylethylenediamine (TEMED) was sourced from Bio-Rad Laboratories.

**Table 1.**
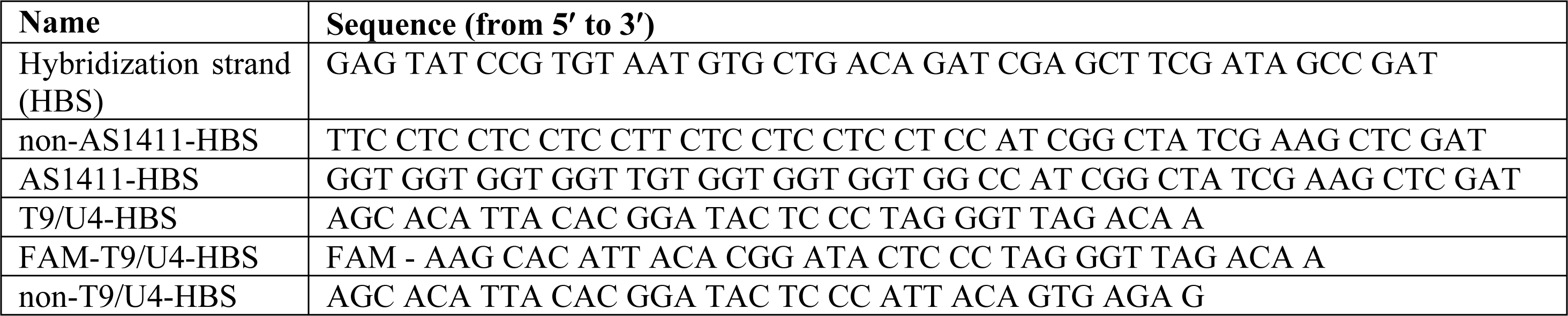
The DNA sequence used in this study.

### Preparation of a molecular hybrid comprising the AS1411 aptamer and T9/U4 ASO

To form a molecular hybrid, T9/U4-HBS, AS1411-HBS, and a hybridization strand with designated complementary sequences were combined in a PBS solution at a final concentration of 10 µM. The mixture was then incubated for 24 hours at room temperature to facilitate the hybridization of these three sequences. This resulting molecular hybrid was designated as AS-T9/U4_MH and confirmed through polyacrylamide gel electrophoresis. Additionally, two control molecular hybrids were prepared by substituting AS1411 aptamer and T9/U4 ASO with two non-specific sequences, denoted as nonAS-T9/U4_MH and AS-nonT9/U4_MH, respectively.

### Evaluation of Stability and Binding of MH in vitro

The T9/U4-HBS sequences were labeled with carboxyfluorescein (FAM) at the 5’ end prior to the formation of MH, allowing for the detection of fluorescence signals under a microscope and flow cytometer. The receptor-cellular binding of MH, mediated by the AS1411 aptamer, was assessed through microscopy and flow cytometry.

#### Microscopic Analysis

SW480, Caco-2 (cancer cells), and CCD841 CoN (normal cells) were seeded at a density of 3×10^5^ cells per well in 96-well plates and incubated for 24 hours. Subsequently, the cells were treated with 10 µM of nonAS-T9/U4_MH and AS-T9/U4_MH for 1.5 hours. After treatment, the cells were washed twice with PBS and stained with 4’,6’-diamidino-2-phenylidoledihydrochloride (DAPI) to visualize nuclei. The cells were then imaged using a fluorescence microscope (Nikon Eclipse Ts2R).

#### Flow Cytometry

SW480, Caco-2, and CCD841 CoN cells (3×10^6^ cells per well) were seeded into 12-well plates and incubated for 24 hours. Afterward, the cells were treated with 5 µM of designated molecules, washed twice with PBS, trypsinized, collected, washed again with PBS, and resuspended in 0.5 mL PBS for binding analysis using flow cytometry.

### Cell Proliferation Assay

To assess cell proliferation, SW480, Caco-2, and CCD841 CoN cells were seeded onto 96-well plates at a density of 5×10^3^ cells per well and incubated for 24 hours. Subsequently, they were treated with various formulations including nonAS-T9/U4_MH, AS-nonT9/U4_MH, and AS-T9/U4_MH for 48 hours. Following treatment, MTS reagent (Celltiter 96®, Promega) was added, and the plates were further incubated for 1 hour before measuring absorbance at 490 nm using a microplate reader (Thermo Scientific, USA).

### Intercalation of Dox into MH

Incorporation of Dox into MH was achieved by incubating 10 µM of MHs with 0.95 µM of Dox at room temperature in the dark for 1.5 hours. The fluorescence intensity of Dox served as an indicator for intercalation, measured using a Varioskan microplate reader (Thermo Scientific, USA) with excitation wavelength set at 480 nm and emission wavelengths recorded from 500 to 800 nm. Dox-incorporated MHs were designated as Dox-loaded AS-T9/U4_MH, Dox-loaded AS-nonT9/U4_MH, and Dox-loaded nonAS-T9/U4_MH.

### Cell Culture

Human colorectal adenocarcinoma cell lines (SW480 and Caco-2) and human normal colon cells (CCD 841 CoN) were obtained from the American Type Culture Collection (ATCC, USA) and cultured in DMEM medium supplemented with 10% fetal bovine serum, 1% penicillin-streptomycin at 37°C in a 5% CO_2_ atmosphere. Additionally, the medium for Caco-2 cells was supplemented with non-essential amino acids. Cells were sub-cultured upon reaching 70–80% confluence.

### Effect of Dox-loaded AS-T9/U4_MH on Cell Proliferation

The effect of Dox-loaded AS-T9/U4_MH on the proliferation of SW480, Caco-2, and CCD841 CoN cells was investigated. Cells were seeded at a density of 5×10^3^ cells per well in 96-well plates, cultured for 24 hours, and then treated with Dox, Dox-loaded nonAS-T9/U4_MH, Dox-loaded AS-nonT9/U4_MH, and Dox-loaded AS-T9/U4_MH, maintaining a concentration of 0.95 µM for Dox and 10 µM for MH. Cell proliferation was determined using the MTS assay.

### Cell Apoptosis

SW480 cells were seeded in 12-well plates at a density of 3×10^6^ cells per well and incubated for 24 hours.

Subsequently, cells were treated with Dox-loaded AS-T9/U4_MH, Dox-loaded AS-nonT9/U4_MH, AS-T9/U4_MH, AS-nonT9/U4_MH, and Dox for 48 hours at a final concentration of 0.95 µM for Dox and 10 µM for MHs. After treatment, cells were collected and subjected to apoptosis analysis using Annexin V-FITC/PI dual staining kit followed by flow cytometry.

### Western blot analysis

The cancer cells (SW480) were treated with 0.95 μM of free Dox, 10 μM of MH formulations: Dox-loaded AS-T9/U4_MH, Dox-loaded AS-nonT9/U4_MH, AS-T9/U4_MH, and AS-nonT9/U4_MH. After 48 hours of incubation, the cells were collected and lysed in RIPA buffer containing protease/phosphatase inhibitor on ice. Quantification of protein concentration was processed using the BCA protein assay (Thermo fisher). A total protein at concentration per lane at 50 μg was loaded into the 10% SDS-PAGE and then transferred to PVDF membranes. Then, the membrane was incubated with blocking buffer (Intercept (TBS), Li-COR) at room temperature for 1 hour and then incubated overnight at 4 °C with the following primary antibodies: hTERT (cat. no. ab32020; 1:1000; 127 kDa; Abcam), vimentin (cat. no. ab92547; 1:1000; 54kDa; Abcam), *β* - actin (1:1000; 42 kDa, Cell signaling), Bcl–2 (1:1000; 25 – 28 kDa, Cell signaling), and Bax (1:1000; 20 kDa, Cell signaling). After incubation, the membrane was washed by 1% TBS-Tween-20. The second antibody, goat antirabbit IgG H&L/HRP antibody (AB6721, Abcam) were incubated on membranes (1:15000) for 1 hour at room temperature and then washed with 1% TBS-Tween-20. Subsequently, all bands were visualized using Odyssey XF imaging system (LI-COR). Densitometric analysis was used to evaluate the protein level, using β-actin as internal control.

### Statistical Analysis

Statistical analysis was conducted using a one-way analysis of variance (ANOVA) test with a minimum of three experimental replicates to ensure statistical power. Results were presented as mean ± standard deviation (SD), with P < 0.05 considered statistically significant.

## Results and Discussion

### Formation of AS-T9/U4 molecular hybrid

To confer the ability to downregulate human telomerase reverse transcriptase (hTERT) expression, we integrated an antisense oligonucleotide known as T9/U4 into our molecular hybrid, referred to as AS-T9/U4_MH. The T9/U4 sequence has been previously documented to effectively reduce hTERT expression [19, 43]. The design of AS1411 aptamer (AS) and T9/U4 antisense oligonucleotide (T9/4U) sequences included additional oligonucleotides to facilitate hybridization with a hybridization strand (HBS). This was achieved by incubating HBS, T9/U4, and AS in a PBS solution at room temperature for 24 hours. The resulting product was characterized using gel electrophoresis. The gel image (Fig 1) demonstrated the successful formation of AS-T9/U4_MH. A distinct band approximately 90 base pairs in size indicated the presence of the molecular hybrid, as its size exceeded that of each individual building block: AS1411 aptamer, T9/U4, and HBS. Rotkrua et al. utilized a similar hybridization technique to form Chol-aptamer molecular hybrid (CAH) with high yield, as reported in their study [44].

**Fig 1.**
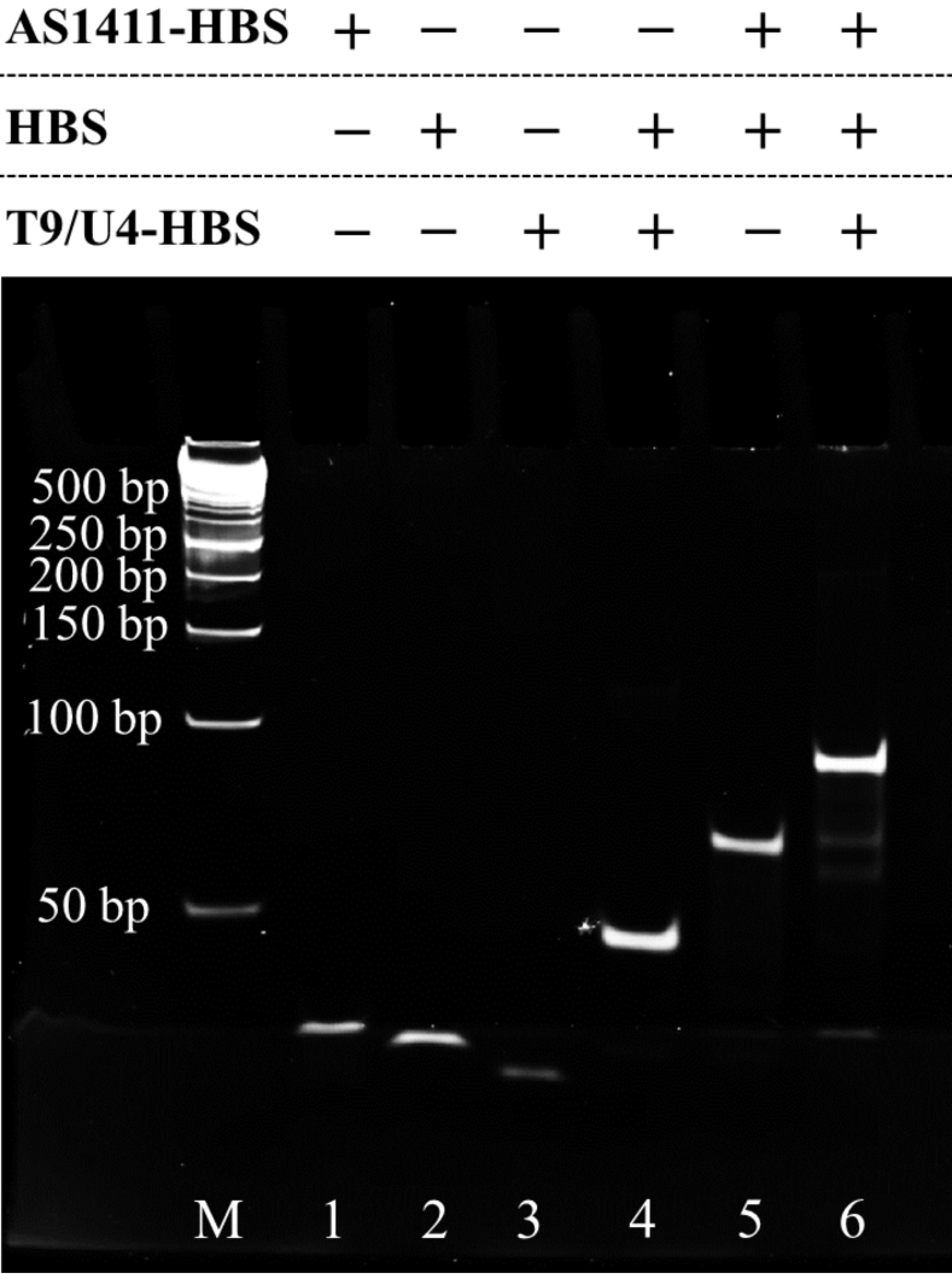
Gel electrophoresis shows the assembly of AS-T9/U4_MH (lane M: DNA marker, lane 1: AS1411-HBS, lane 2: HBS, lane 3: T9/U4-HBS, lane 4: HBS + T9/U4-HBS, lane 5: HBS + AS-HBS, and lane 6: HBS + T9/U4-HBS + AS-HBS).

### Stability and binding of MH *in vitro*

#### Fluorescence microscope

To demonstrate the potential of AS1411 aptamer as a carrier for T9/U4 to target tumor tissue, we utilized fluorescence microscopy to observe Caco-2, SW480, and CCD841 CoN cells after treating them with FAM-labeled MH (Fig 2). The fluorescence images revealed that FAM-labeled AS-T9/U4_MH specifically bound to SW480 cells. This specificity can be attributed to the known ability of AS1411 aptamer to bind to nucleolin, which is overexpressed on the surface of SW480 cells, as reported in the literature [44]. Conversely, the fluorescence images of Caco-2 and CCD841 CoN cells exhibited lower intensity, suggesting less binding of the FAM-labeled AS-T9/U4_MH. This discrepancy may arise from the lower availability of nucleolin in Caco-2 and CCD841 cells. Research by Dean and Kenny has shown that the expression level of native nucleolin on the cell surface of intestinal Caco-2 cells is notably low, a finding corroborated by the detectability of this molecule using nucleolin antibodies such as MS-3 from Santa Cruz Biotechnology and ab22758 from Abcam [45, 46]. Similarly, Lohlamoh et al. found that nucleolin mRNA expression in SW480 cells was significantly higher than in CCD841 CoN cells [47]. Furthermore, Duncan et al. determined nucleolin’s overexpression in both cancer cells and normal cells, including HS-27 (skin fibroblast), WI-38 (lung fibroblast), and MCF-10A (epithelial mammary cell). Their study highlighted that the nucleolin level in normal fibroblast cells was approximately four times lower than in fibroblast-like cancer cells such as HT-1080 and SK-MEL2 [48]. These findings collectively support the differential binding of AS-T9/U4_MH to tumor cells versus normal cells based on nucleolin expression levels, reinforcing the potential of AS1411 aptamer as a targeted delivery vehicle for T9/U4 to tumor tissues.

**Fig 2.**
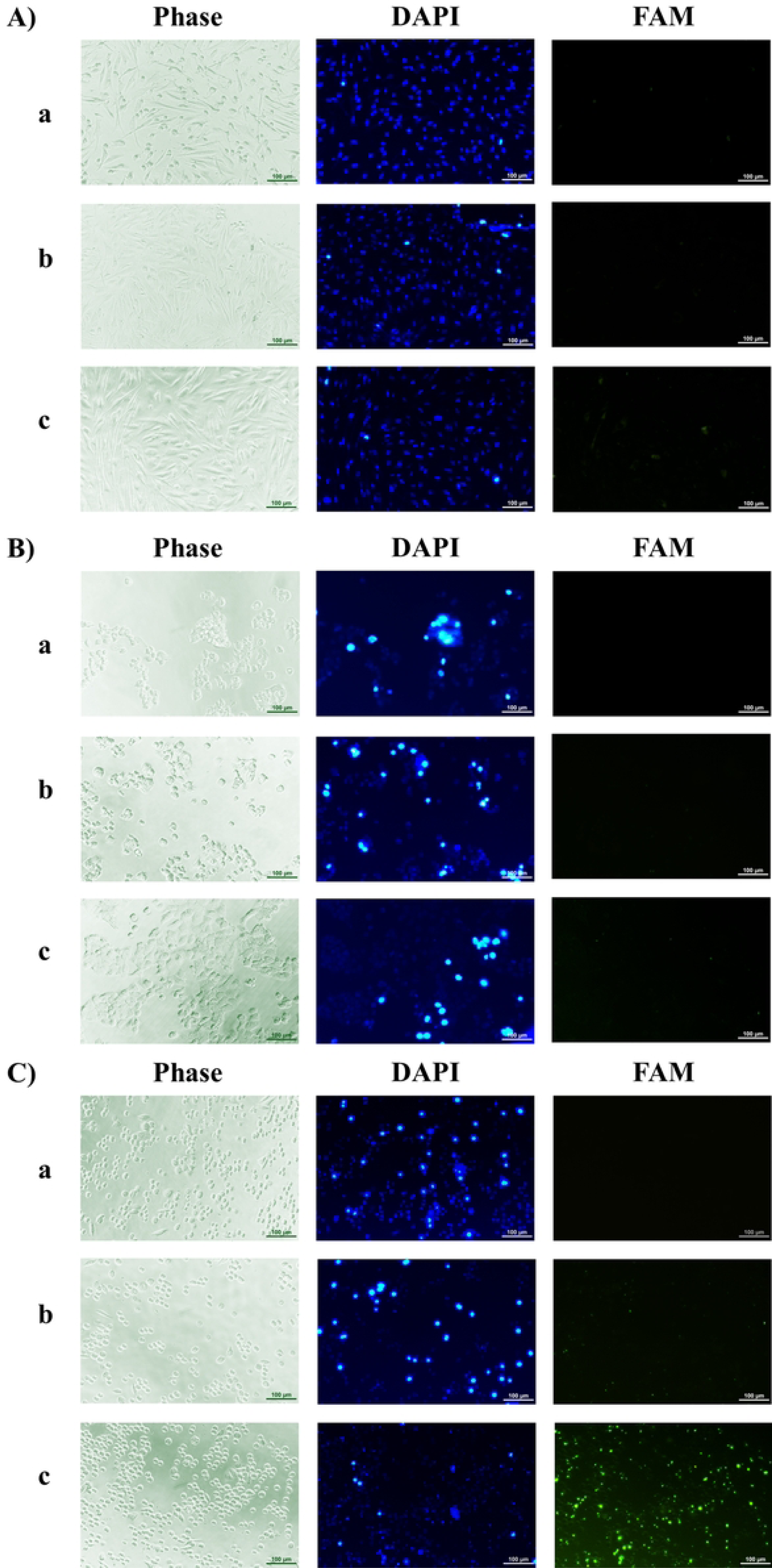
Fluorescence image of CCD841 CoN (A), Caco-2 (B), and SW480 (C), treated with (a) no treatment, (b) FAM-labeled nonAS-T9/U4_MH, and (c) FAM-labeled AS-T9/U4_MH.

#### Flow cytometry

Flow cytometry analysis was employed to evaluate the intracellular fluorescent intensity subsequent to the treatment of SW480, Caco-2, and CCD841 CoN cells with FAM-labeled nonAS-T9/U4_MH and FAM-labeled AS-T9/U4_MH. AS-T9/U4_MH treatment notably resulted in a significantly detected fluorescent intensity in SW480 cells compared to other molecular hybrids (Figs 3A and 3B). Conversely, Caco-2 and CCD841 CoN cells exhibited minimal fluorescence signals when exposed to the MH formulation utilized for cell treatment. These observations are consistent with our microscopy assay findings, further corroborating the preferential binding and uptake of AS-T9/U4_MH by SW480 cells in comparison to other cell types.

**Fig 3.**
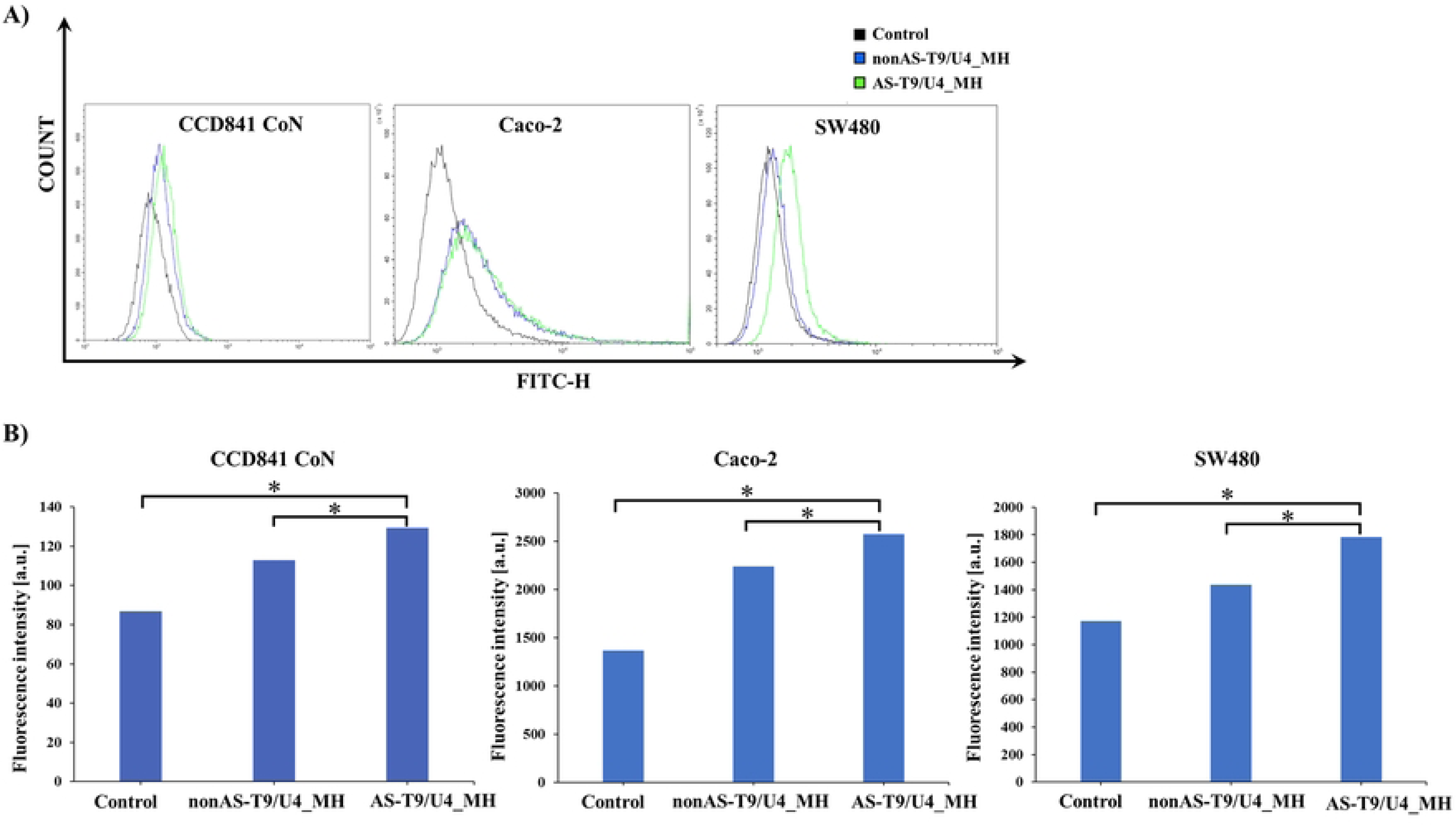
AS-T9/U4_MH showed specific binding to SW480 cells. A, B) Flow cytometry histogram of CCD841 CoN (left), Caco-2 (middle) and SW480 (right). The black line represents cells without MH treatment. The blue line represents cells treated with FAM-labeled nonAS-T9/U4_MH. The green line represents cells treated with FAM-labeled AS-T9/U4_MH. The cells were treated at 37 °C for 1.5 h, and no treatment as a control. *P < 0.05. The data are presented as means ± SD, n = 3.

### Cell proliferation assay

To evaluate the effect of AS-T9/U4_MH on cell proliferation, we conducted MTS assays using SW480, Caco-2, and CCD841 CoN cells. The results revealed that AS-T9/U4_MH exerted an anti-proliferative effect specifically on SW480 cells, while demonstrating no significant effect on Caco-2 and CCD841 CoN cells (Fig 4). Notably, the cell viability of SW480 cells treated with AS-T9/U4_MH was lower compared to those treated with nonAS-T9/U4_MH. This specificity can be attributed to the presence of AS1411 aptamer within the molecular hybrid, as the aptamer is capable of recognizing nucleolin molecules present on the surface membrane of SW480 cells [49]. Research by Emilio et al. further supports the specificity of AS1411 aptamer, demonstrating its ability to significantly inhibit the phosphorylation of nucleolin in human umbilical vein endothelial cells (HUVEC), whereas control formulations such as scramble sequences and ranibizumab had no effect on nucleolin phosphorylation [50]. Conversely, AS-T9/U4_MH exhibited no discernible effect on Caco-2 and CCD841 CoN cells, likely due to their lower nucleolin expression levels [48]. These findings underscore the targeted anti-proliferative effect of AS-T9/U4_MH on SW480 cells, driven by the specific interaction between AS1411 aptamer and nucleolin, highlighting its potential as a targeted therapeutic agent for tumors with elevated nucleolin expression levels.

**Fig 4.**
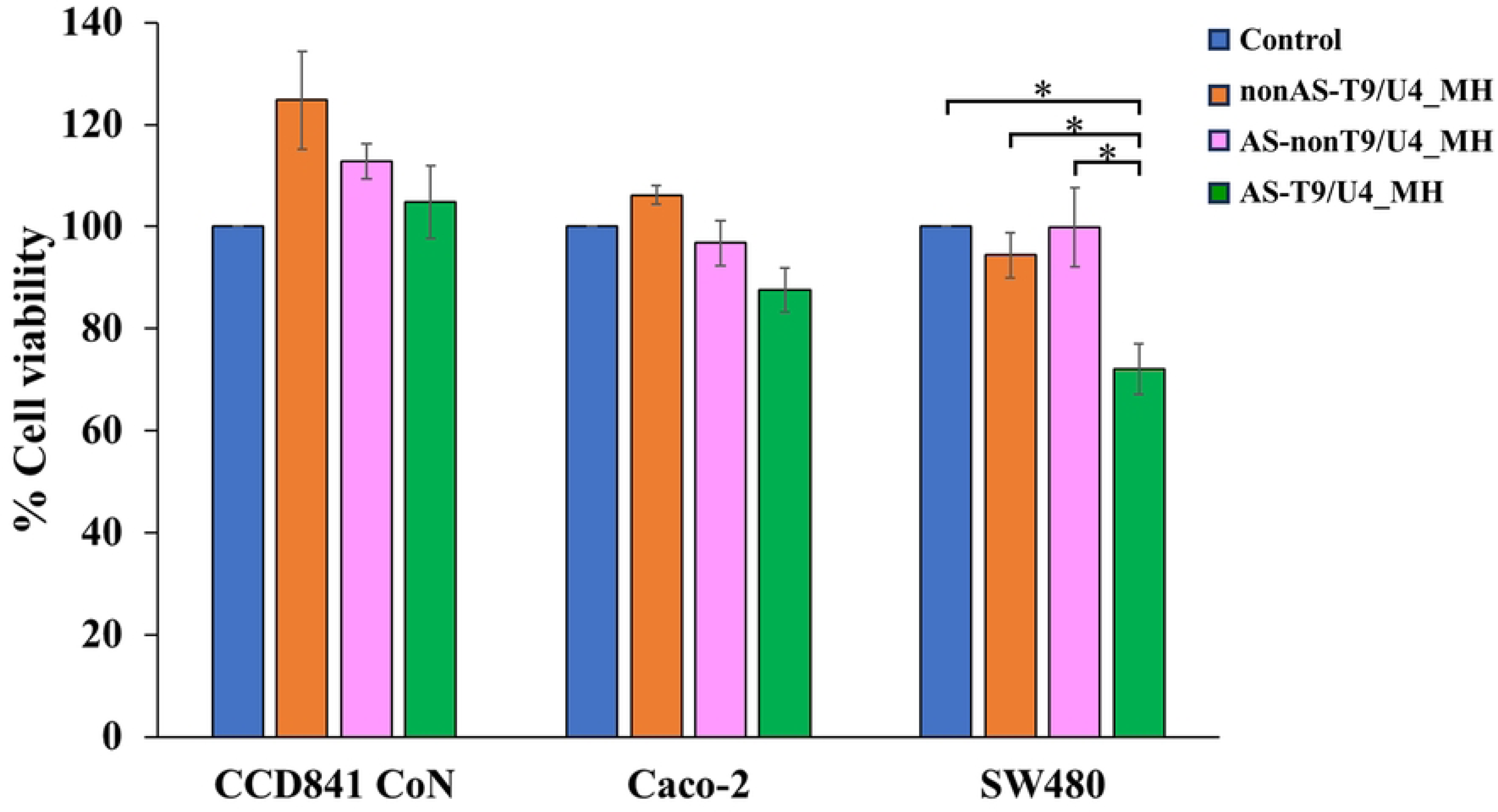
The cell viability of CCD841 CoN, Caco-2, and SW480 cells after treatments at 48 h of 10 μM nonAS-T9/U4_MH, AS-nonT9/U4_MH, AS-T9/U4_MH and no treatment as a control. The values are presented as means ± SD, n = 3, \**P* < 0.05

### Intercalation of Dox into AS-T9/U5_MH

Fluorescence spectroscopy served as the method of choice to assess the intercalation of Dox into MH by detecting fluorescence quenching. In this work, Dox was excited using a light source at 480 nm, and the emitted fluorescence was detected with a maximum signal at 590 nm (Fig 5). The observed fluorescence quenching of Dox upon its incorporation into the molecular hybrids indicates the complete intercalation of Dox within the double helix of DNA [51]. This phenomenon confirms the successful integration of Dox into the MH structure, validating its potential for targeted drug delivery.

**Fig 5.**
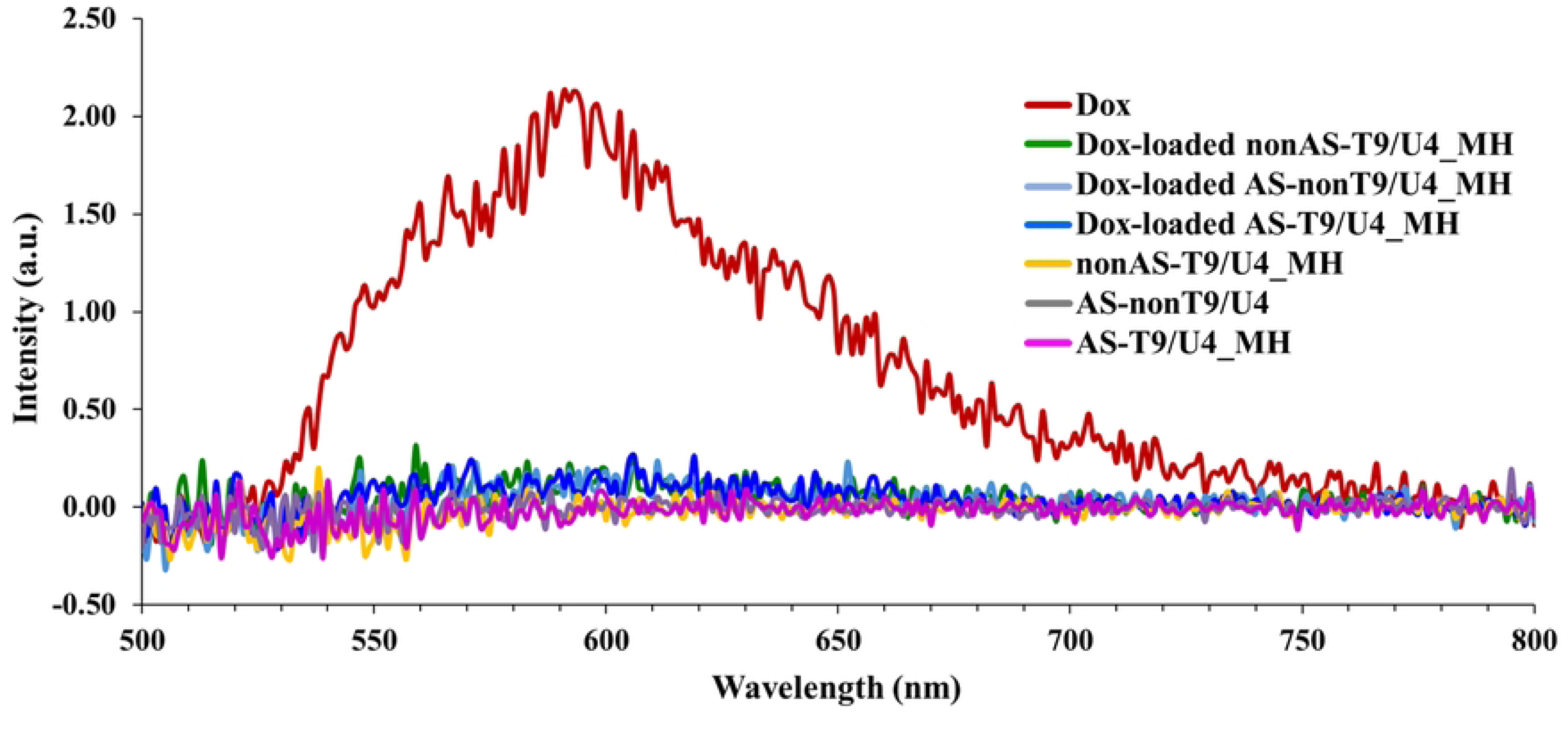
Fluorescence spectra of 0.95 μM of Dox and 10 μM of Dox-loaded nonAS-T9/U4_MH, Dox-loaded AS-nonT9/U4_MH, Dox-loaded AS-T9/U4_MH, nonAS-T9/U4_MH, AS-nonT9/U4_MH, and AS-T9/U4_MH.

### Effect of Dox-loaded AS/T9/U4_MH on cell proliferation

Free Dox demonstrated toxicity towards Caco-2 and SW480 cells, while showing no impact on CCD841 CoN cells (Fig 6), likely due to its therapeutic window being greater compared to Caco-2 and SW480 cells [44]. In the context of Caco-2 cells, upon intercalation of Dox into the DNA duplex of our molecular hybrids, a significant reduction in cytotoxicity was observed. This reduction can be attributed to the DNA duplex system’s effectiveness in minimizing the cytotoxic effects of this anti-cancer drug, as previously reported in our study [44]. For SW480 cells, Dox-loaded AS-T9/U4_MH demonstrated the specific delivery of Dox to the cells when compared to the other two control molecular hybrids, as evidenced by the cell viability results. The enhanced effectiveness of this MH can be attributed to the synergistic effect resulting from the presence of both Dox and T9/U4 ASO. This finding aligns with previous studies in the literature. For instance, Abaza et al. demonstrated that the antiproliferative effects of colorectal cancer cells were synergistically enhanced by combining c-myc antisense phosphorothioate oligonucleotides with anticancer drugs such as taxol, 5-FU, Dox, and vinblastine [52]. Similarly, Jhaveri et al. showed that combining human α isoform folate receptor (αhFR) antisense oligonucleotides with Dox resulted in a five-fold reduction in αhFR expression in breast cancer cells [53]. The specificity of Dox-loaded AS-T9/U4_MH was achieved through the presence of AS1411 aptamer, as Caco-2 and CCD841 CoN cells have lower availability of nucleolin [45, 48]. Consistent with previous research, Dox intercalated into MH exhibited reduced toxicity compared to free Dox, suggesting that the MH strategy may offer a means to mitigate the adverse effects of Dox [44]. Thus MH might be a promising approach for targeted and less toxic drug delivery in cancer treatment.

**Fig 6.**
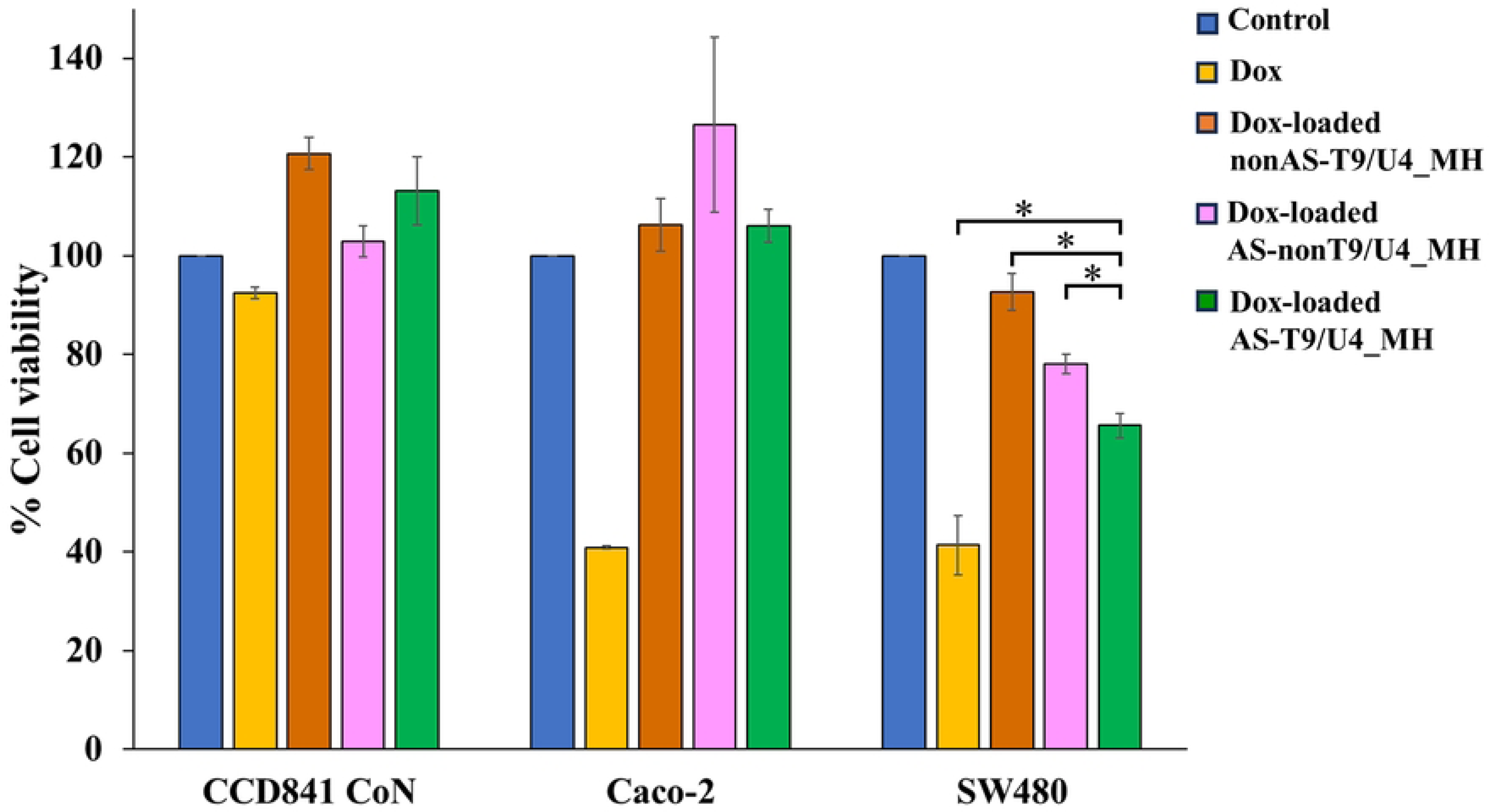
The cell viability of CCD841 CoN, Caco-2, and SW480 cells after treatments at 48 h of 0.95 μM Dox, 10 μM Dox-loaded nonAS-T9/U4_MH, Dox-load AS-nonT9/U4_MH, Dox-load AS-T9/U4_MH, and no treatment as a control. The values are presented as means ±SD, n = 3, \**P* < 0.05.

### Effect of Dox-loaded AS/T9/U4_MH on Cell apoptosis

Flow cytometry was employed to investigate the impact of T9/U4 ASO and Dox-loaded MHs on apoptosis in SW480 cells. The results revealed that the percentage of cell apoptosis was significantly higher in cells treated with AS-T9/U4_MH compared to those treated with AS-nonT9/U4_MH. This observation suggests that T9/U4 ASO effectively inhibited SW480 cell proliferation in vitro (Fig 7). Furthermore, the incorporation of Dox into AS-T9/U4_MH resulted in a more pronounced effect on apoptosis compared to Dox-loaded AS-nonT9/U4_MH, indicating a synergistic effect. This enhanced efficacy of Dox-loaded AS-T9/U4_MH indicates the potential of MHs as effective carriers for delivering therapeutic agents, such as Dox, in combination with ASOs for targeted cancer therapy. These findings show the promising synergistic effects of combining ASO-mediated inhibition of cell proliferation with the cytotoxic effects of Dox, offering new insights into potential strategies for enhancing cancer treatment efficacy.

**Fig 7.**
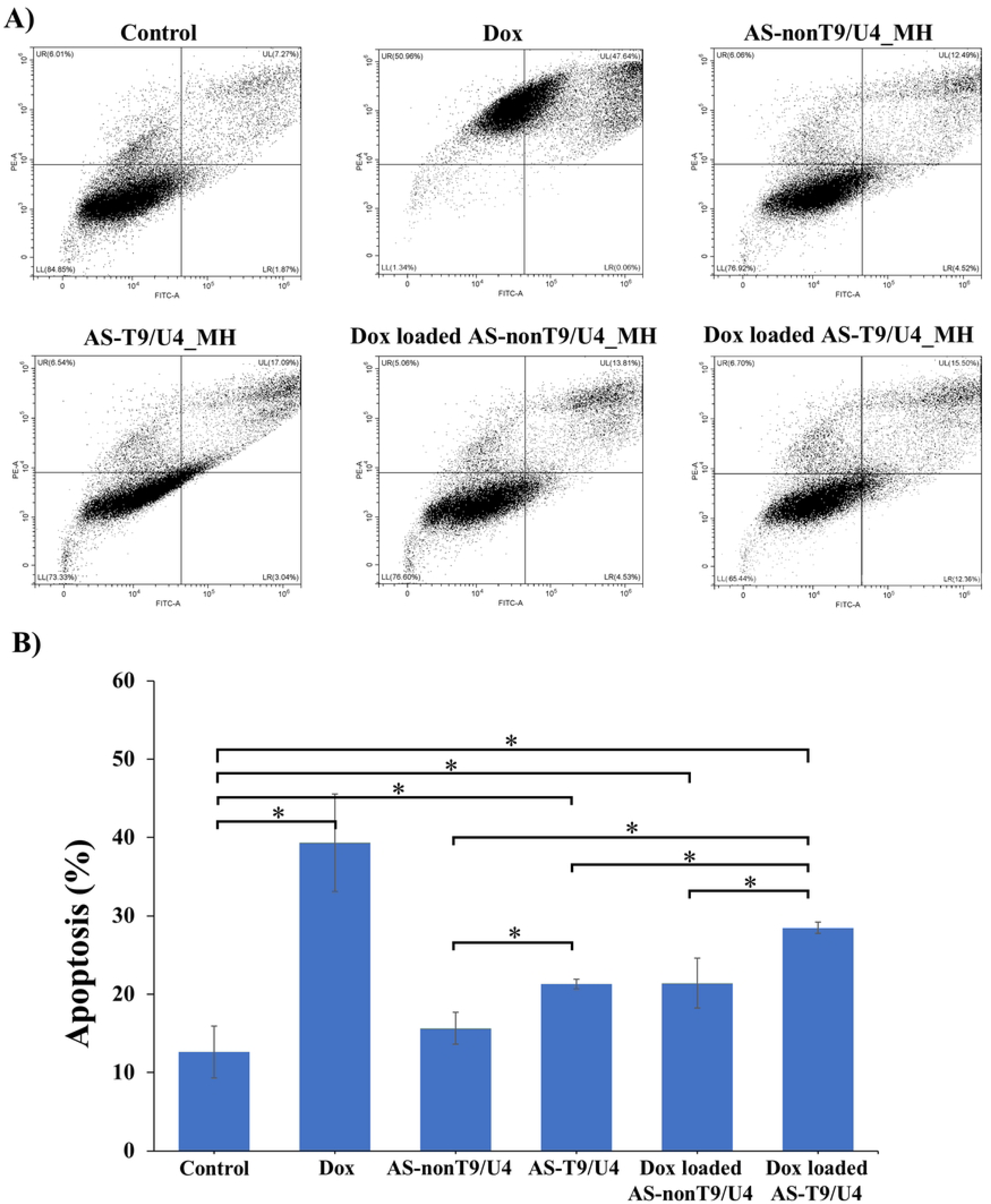
Cell apoptosis of SW480 cells treated with 0.95 μM of Dox and 10 μM of AS-T9/U4_MH, AS-nonT9/U4_MH, Dox-loaded AS-T9/U4_MH, and Dox-loaded AS-nonT9/U4_MH at 48 h, and no treatment as a control. A) flow cytometry plot. B) Cell apoptosis count. The values are presented as means ±SD, n = 3, *P < 0.05.

### Effect of Dox-loaded AS/T9/U4_MH on hTERT and vimentin expression

The expression of hTERT and vimentin plays a crucial role in a process of epithelial-mesenchymal transition (EMT), which is pivotal in cancer progression by transforming epithelial cells into a mesenchymal state, thus significantly contributing to tumor development [54–56]. To explore how T9/U4 ASO regulates hTERT expression, we evaluated the hTERT levels in SW480 cells. Decreased hTERT levels led to the suppression of telomerase activity, which in turn benefitted by reducing cancer cell proliferation [57], enhancing apoptosis [58], and improving sensitivity to chemotherapy [59]. SW480 cells treated with AS-T9/U4_MH and Dox-loaded AS-T9/U4_MH showed a significant reduction in hTERT expression levels compared to cells treated with MHs lacking ASO (Figs 8A and 8B). These results suggested that T9/U4 ASO effectively downregulated hTERT levels in SW480 cells. Our results aligned with several studies indicating that T9/U4 modified with phosphorothioate (PS) had a potent effect on inhibiting telomerase activity in HL-60, A549-luc, and U-251 MG cells [43, 60]. Moreover, the AS1411 aptamer and Dox in the MH contributed to lowering hTERT expression. The AS1411 aptamer naturally forms a G-quadruplex structure, effectively inhibiting telomerase activity by targeting telomeric G-quadruplexes and stabilizing those present in the hTERT promoter [61]. Dox was also reported to inhibit telomerase activity in SW480 cells [62], exhibiting a toxic effect that reduces both telomerase activity and hTERT expression [63]. Thus, loading Dox into AS-T9/U4_MH could further decrease hTERT level. These results indicated that T9/U4 ASO, AS1411 aptamer, and Dox acted synergistically to lower hTERT expression.

**Fig 8.**
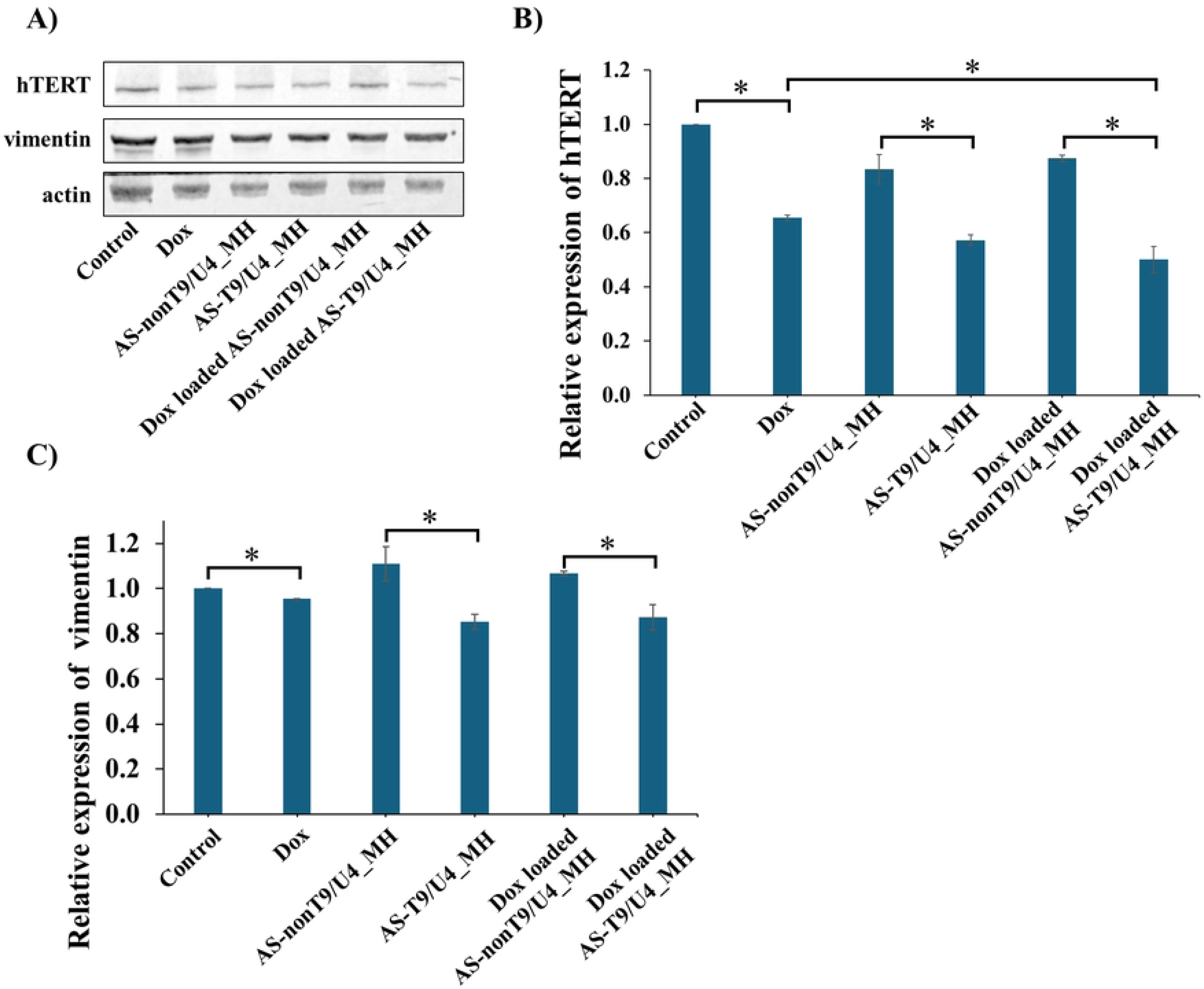
(A) Western blot analysis of hTERT and vimentin expression in SW480 cells treated with Dox, AS-T9/U4_MH, AS-nonT9/U4_MH, Dox-loaded AS-T9/U4_MH, and Dox-loaded AS-nonT9/U4_MH, and no treatment as a control. The relative expression of (B) hTERT and (C) vimentin. The values are presented as means ±SD, n = 3, \**P* < 0.05.

The vimentin expression level was further investigated. Treatment with both AS-T9/U4_MH and Dox-loaded AS-T9/U4_MH led to a decrease in vimentin expression in SW480 cells compared to cells treated with MHs lacking the specific ASO (Figs 8A and 8C). These findings suggested that T9/U4 ASO effectively reduced vimentin levels through hTERT dysfunction. Additionally, the results indicated that Dox and the AS1411 aptamer slightly decreased vimentin expression in vitro. The reduced effect of Dox on vimentin expression in SW480 cells was also noted in a study on MFT-16 cells, which investigated the impact of vimentin on drug resistance. This study revealed that a mutant vimentin attached to mitochondria, enhancing cell membrane potential, consequently resulting in decreased Dox effectiveness [64]. As for the AS1411 aptamer, vimentin expression in SNU-761 hepatocellular carcinoma (HCC) remained unaltered after treatment with the aptamer, likely due to a key survival signaling pathway of HCC involving the PI3K/Akt or ERK1/2-MAPK pathway [65].

### Effect of Dox-loaded AS/T9/U4_MH on Bcl-2 and Bax expression

Bcl-2 and Bax play pivotal roles in apoptotic pathways, with Bcl-2 exhibiting anti-apoptotic properties and Bax known for its pro-apoptotic function [66]. Investigating the impact of our MHs on the expression levels of Bcl-2 and Bax in SW480 cells can provide crucial insights into related apoptotic pathways. Upon treatment with MHs, a notable decrease in Bcl-2 expression levels was observed in cells treated with free Dox, AS-T9/U4_MH, Dox-loaded AS-nonT9/U4_MH, and Dox-loaded AS-T9/U4_MH formulations (Figs 9A and 9B). Notably, the MH formulation containing Dox, the aptamer, and the targeted ASO exhibited the most significant reduction in Bcl-2 expression. Dox facilitated apoptosis by upregulating the expression of Bax, caspase-8, and caspase-3, while concurrently downregulating Bcl-2 expression, as demonstrated in MCF-10F, MCF-7, and MDA-MB-231 breast cancer cell lines [67]. Interestingly, the AS1411 aptamer showed no effect on Bcl-2 expression, despite its reported inhibition of SW480 cell proliferation cells [47]. Strikingly, T9/U4 ASO markedly influenced Bcl-2 expression, emphasizing its efficacy in promoting cell apoptosis. Regarding Bax protein, the MH containing Dox and T9/U4 ASO notably increased its expression level (Figs 9A and 9C). Previous literature suggested that Dox induces apoptosis in MCF-7 breast cancer cells via the mitochondrial pathway by decreasing Bcl-xL expression and increasing Bax expression in a dose-dependent manner [68]. The expression patterns of both proteins indicated that Dox-loaded AS-T9/U4_MH affected this apoptotic pathway, with Dox and the ASO synergistically enhancing apoptosis in SW480 cells.

**Fig 9.**
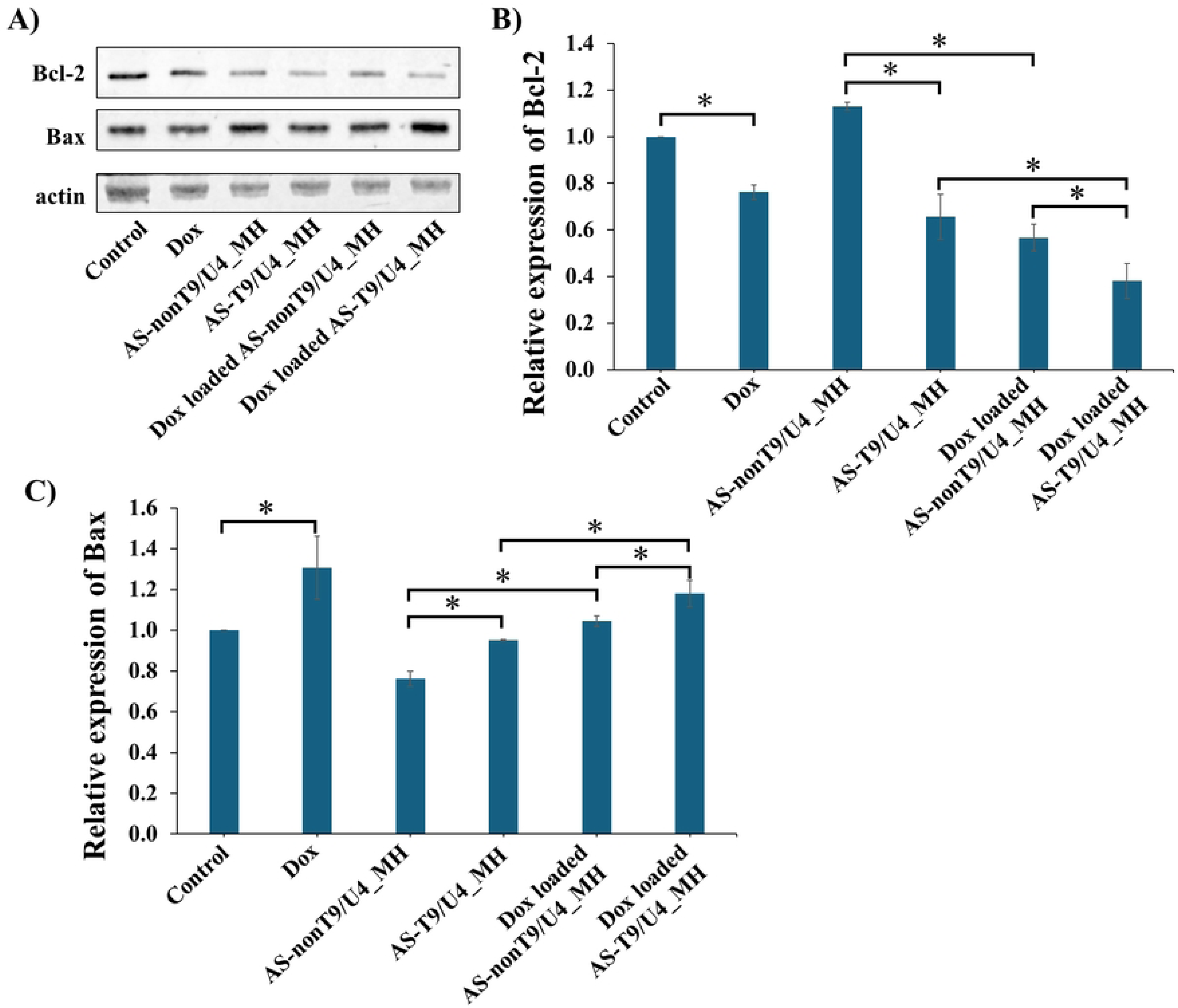
(A) Western blot analysis of Bcl-2 and Bax expression in SW480 cells treated with Dox, AS-T9/U4_MH, AS-nonT9/U4_MH, Dox-loaded AS-T9/U4_MH, and Dox-loaded AS-nonT9/U4_MH, and no treatment as a control. The relative expression of (B) Bcl-2 and (C) Bax. The values are presented as means ±SD, n = 3, \**P* < 0.05.

## Conclusions

This work accomplished following aspects. The following achievements were made in this study. Gel electrophoresis confirmed the formation of MH, indicating successful preparation of AS–T9/U4_MH. The MH created was found to specifically target SW480 cells, as demonstrated by cell viability assays, fluorescence microscopy, and flow cytometry, while showing no significant impact on Caco-2 and CCD841 CoN cells. AS – T9/U4_MH effectively served as a carrier for delivering ASO and Dox, leading to a synergistic effect observed in cell viability results. Interestingly, loading Dox into the DNA double helix resulted in reduced cytotoxicity. Furthermore, Dox-loaded AS-T9/U4_MH effectively downregulated hTERT and vimentin expression in SW480 cells, indicating its potential in inhibiting epithelial-mesenchymal transition (EMT) and tumor progression. Lastly, the impact of MHs on apoptotic pathways was investigated, revealing a significant decrease in Bcl-2 expression and increased Bax expression in SW480 cells treated with Dox-loaded AS-T9/U4_MH. This suggests the involvement of the mitochondrial pathway in apoptosis induction, with synergistic effects between Dox and T9/U4 ASO. These findings demonstrate the potential of oligonucleotide hybridization in forming versatile macromolecular complexes with multiple functionalities. The success of this approach suggests that MH could serve as efficient drug delivery systems for cancer therapy.

## Acknowledgment

This study was supported by Thailand Science Research and Innovation Fundamental Fund (Contract No. TUFF 27/2567), and Thammasat University Research Unit in Innovation of Molecular Hybrid for Biomedical Application. In addition, BS was financially supported for the travelling, and the participation of JCA 82^nd^ annual meeting by Thammasat University and the Japanese Cancer Association. KJ received scholarship from Faculty of Science and Technology, Thammasat University and National Nanotechnology Center (NANOTEC), National Science and Technology Development Agency (NSTDA).

